# Low Quantity single strand CAGE (LQ-ssCAGE) maps regulatory enhancers and promoters

**DOI:** 10.1101/2020.08.04.231969

**Authors:** Hazuki Takahashi, Hiromi Nishiyori-Sueki, Jordan A. Ramilowski, Masayoshi Itoh, Piero Carninci

**Affiliations:** RIKEN Center for Integrative Medical Sciences (IMS), Yokohama, Kanagawa 230-0045, Japan; RIKEN Preventive Medicine and Diagnosis Innovation Program (PMI), Saitama 351- 0198, Japan

**Keywords:** Cap Analysis of Gene Expression (CAGE), High throughput CAGE library preparation, Promoter, Enhancer, eRNA, TSSs

## Abstract

The Cap Analysis of Gene Expression (CAGE) is a powerful method to identify the Transcription Start Sites (TSSs) of capped RNAs while simultaneously measuring transcripts expression level. CAGE allows mapping at single nucleotide resolution at all active promoters and enhancers. Large CAGE datasets have been produced over the years from individual laboratories and consortia, including the Encyclopedia of DNA Elements (ENCODE) and Functional Annotation Of the Mammalian Genome (FANTOM). These datasets constitute open resource for TSS annotations and gene expression analysis. Here we provide an experimental protocol for the most recent CAGE method called Low Quantity (LQ) single strand (ss) CAGE “LQ-ssCAGE”, which enables cost-effective profiling of low quantity RNA samples. LQ-ssCAGE is especially useful for samples derived from cells cultured in small volumes, cellular compartments such as nuclear RNAs or for samples from developmental stages. We demonstrate the reproducibility and effectiveness of the method by constructing 240 LQ-ssCAGE libraries from 50 ng of THP-1 cell extracted RNAs and discover lowly expressed novel enhancer and promoter-derived lncRNAs.

## 1. Introduction

Cap Analysis of Gene Expression (CAGE) is a method to profile RNA expression and precisely identify promoters and regulatory elements such as enhancers. CAGE can identify not only protein coding RNAs but also non-coding RNAs, which are capped when produced by RNA polymerase II. CAGE technology has been developed and modified over the years to follow advanced sequencing technologies and biological interests [1–6].

Classically, promoters and enhancers are defined as genomic elements which proximally initiate and distally enhance transcription, respectively. A part of promoters is highly cell type specific and dynamically associate with nuclear architecture such as histone modifications and nucleosome depleted regions, together with the enhancer (e) RNAs. Unlike RNA derived from standard promoters, eRNAs are typically unstable and poorly adenylated transcripts, and located up to 1 Mb apart from the core promoter in the nucleus. However, many transcriptome analyses have shown that distal enhancers might also play roles into the promoter activity (reviewed in [7]). We previously analyzed cell nuclear comprehensive CAGE dataset from polyadenylated (poly-A) and non-poly-A RNAs with chromatin architectural datasets from ENCODE consortium, which showed that promoters are associated with the complex three-dimensional interconnected chromatin network [8]. Enhancers and promoters are known to act as highly cell type specific regulatory elements [8–10] and active enhancers are likely to transcribe eRNAs that are interacting through chromatin looping with promoters [11, 12]. Further analysis of enhancer – promoter (EP) interactions of ENCODE CAGE dataset also showed that eRNA expressions associated with predicted EP interactions are clearly cell type specific [12].

Importantly, together with eRNAs and promoters, long noncoding RNAs (lncRNAs) overlap regions known to be involved in human genetic traits. In particular, these elements overlap expression quantitative trait loci (eQTL) and single nucleotide polymorphisms (SNPs) associated with genome-wide association studies (GWAS). In particular, lncRNAs promoters/enhancers are significantly and specifically co-expressed in cell types that are related to each with human disease [13].

In addition, the single-base resolution of CAGE TSS mapping has revealed that transcription initiation at thousands of promoters dynamically shifts throughout the specific zebrafish early developmental stage, which are orchestrated with epigenome modifications [14].

To further broaden these analyses, we have developed the high throughput LQ-ssCAGE method, which based on previously developed cap trapper technology [15]; the method is designed to analyze promoters and enhancers usages with single nucleotide resolution from large number of sample, which may include human patient samples, early developmental stage samples and specific cell types. An initial study using the LQ-ssCAGE method demonstrated that antisense lncRNA-mRNA pairs have specific patterns in the cellular compartment from zebrafish early developmental stage [16].

The LQ-ssCAGE presented here captures both poly-A and non-poly-A transcripts by using a 15 nucleotides (N15) random primer in the reverse transcriptase (RT) reaction, before CAP trapper procedures (*see* the workflow of the protocol in Fig. 1). A key advantage of the LQ-ssCAGE consists in using small quantities of RNA yet avoiding PCR amplification, which would otherwise lead to biased and noisy quantification of expression.

**Fig. 1.**
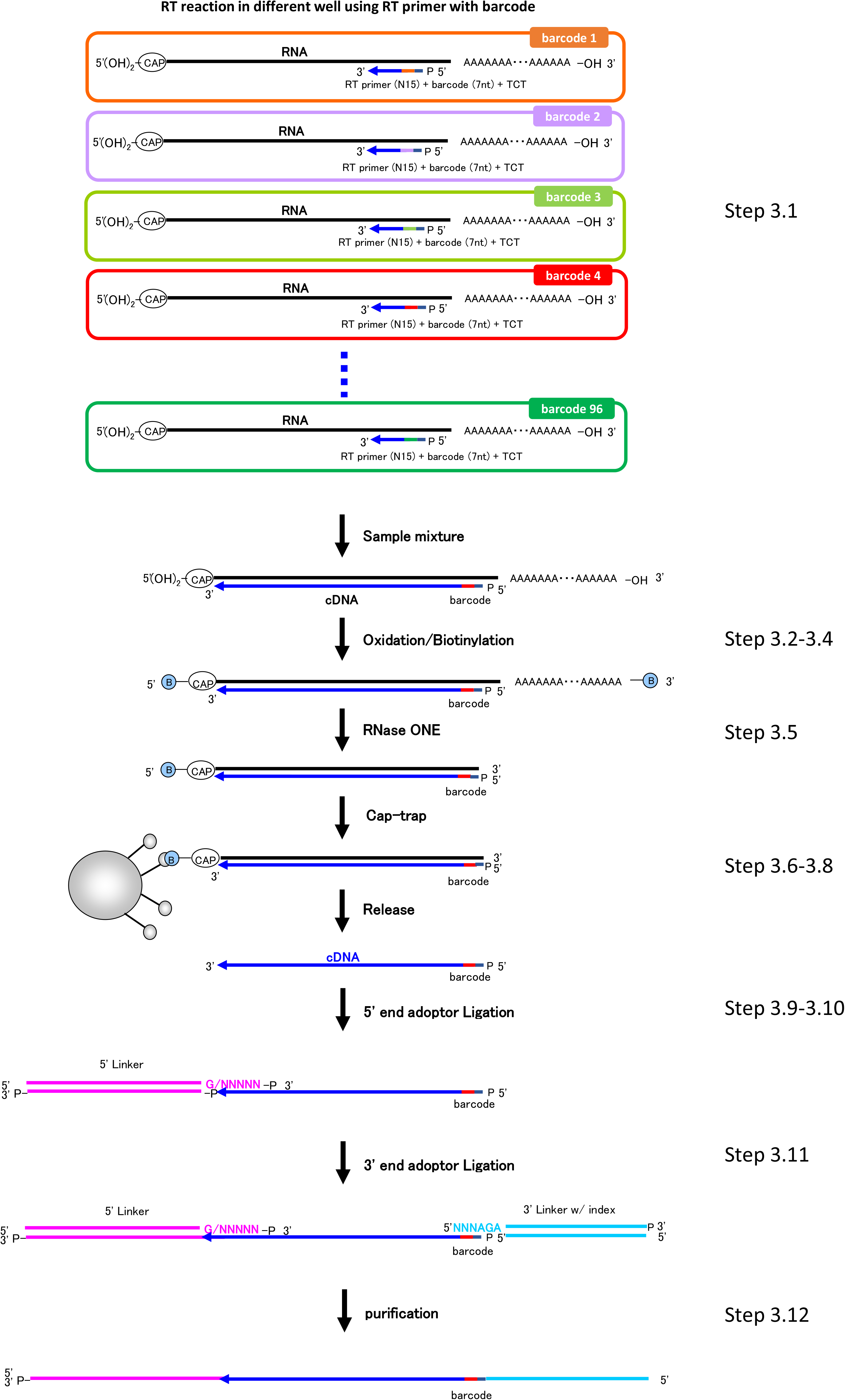
The workflow of LQ-ssCAGE library preparation. The step numbers correspond to those in the methods.

Compared to other CAGE methods [4, 5], the LQ-ssCAGE protocol is further simplified, as the material that is loaded in the sequencer consists simply of single stranded cap selected cDNA. In summary, LQ-ssCAGE protocol (1) can work with RNA amount as little as 25 ng, (2) shortens the preparation time (less than 3 days) and (3) allows for preparing multiple libraries in parallel in microtiter plates. Such libraries can be efficiently sequenced on various Illumina sequencing platforms by using molecular indexes and original barcode identifiers. With a proper organization of libraries preparation workflow, an operator can easily prepare 96 samples simultaneously in a 96-well plate.

To show reproducibility and to confirm that the method allows identification of key regulatory elements, including enhancers and lncRNA promoters, we prepared LQ-ssCAGE libraries from human acute monocytic leukemia (THP-1) cells and analyzed regulatory RNAs comparing our data to FANTOM Cage Associated Transcripts (CAT) gene models and annotations [13]. As examples, we identified enhancers and promoter-derived regulatory RNAs, including GAS5 promoter-derived lncRNA, previously shown to be associated with apoptosis pathway in THP-1 cells [17], and a novel bidirectional region transcribing e-lncRNAs.

## 2. Materials

### 2-1. Equipment

1. 0.2 mL Polypropylene PCR Tube Strips and Domed Cap Strips, 8 Tubes/Strip, 8 Domed Caps/Strip, Clear, Nonsterile (CORNING AXYGEN, special treatment for smooth surface)
2. 1.5 mL Maxymum Recovery Snaplock Microcentrifuge Tube, Polypropylene, Clear, Nonsterile, 250 Tubes/Pack, 10 Packs/Case (CORNING AXYGEN, special treatment for smooth surface)
3. 16 Well Polypropylene PCR Microplate, Clear, Nonsterile (CORNING AXYGEN, special treatment for smooth surface)
4. 96-well Polypropylene PCR Microplate, No Skirt, Clear, Nonsterile (CORNING AXYGEN, special treatment for smooth surface)
5. PCR 1 × 8 Strip Domed Caps, Fit 0.2 mL PCR Tube Strips, Clear, Nonsterile (CORNING AXYGEN, special treatment for smooth surface)
6. X-Pierce Sealing Films, Sterile (EXCEL Scientific, Inc.)
7. Low binding barrier tips of 10 μL (0.1 - 10 μL), 20 μL (1-20 μL), 200 μL (1-200 μL) and 1000 μL (100-1000 μL).
8. PIPETMAN P2, P20, P200 and P1000
9. Thermal Cycler
10. Centrifuge for plate, PCR tube and 1.5 mL tube.
11. 8 channel pipettes for 0.5 - 10 μL and 10 - 100 μL and 30 - 300 μL
12. Vortex mixer
13. miVAC DNA (SP Scientific Genevac)
14. miVac rotor for micro plate (SP Scientific Genevac)
15. Dynabeads MPC-S (Magnetic Particle Concentrator) (Thermo Fisher Scientific)
16. DynaMag-96 Side Skirted Magnet (Thermo Fisher Scientific)

### 2-2. Commercial Reagents

1. Agencourt AMPure XP (BECKMAN COULTER)
2. Agencourt RNAClean XP (BECKMAN COULTER)
3. KAPA Library Quantification Kits (KAPA BIOSYSTEMS)
4. 5 M Sodium Chloride, Molecular Biology Grade
5. RNase One Ribonuclease (Promega)
6. Lithium chloride solution 8 M, for molecular biology, ≥99% (Sigma-Aldrich)
7. Sodium acetate buffer solution BioXtra, pH 7.0±0.05 (25 °C) for molecular biology, 3 M, non-sterile; 0.2 μm filtered (Sigma-Aldrich)
8. Sodium periodate ACS Reagent Grade (Sigma-Aldrich)
9. DNA Ligation Kit Mighty Mix (Takara Bio Inc)
10. Ribonuclease H (RNase H) (20~60 U/μL) (Takara Bio Inc)
11. 10 mM dNTP Mix (Thermo Fisher Scientific)
12. Dynabeads M-270 Streptavidin (Thermo Fisher Scientific)
13. RNase Decontamination Solution
14. SuperScript III Reverse Transcriptase (Thermo Fisher Scientific)
15. UltraPure 0.5 M EDTA (pH 8.0)
16. UltraPure DNase/RNase-Free Distilled Water
17. Biotin (Long Arm) Hydrazide (Vector Laboratories) 18. 0.5 M EDTA (pH 8.0)
18. 10w/v% Polyoxyethylene (20) Sorbitan Monolaurate Solution
19. 1 M Tris-HCl (pH 7.0, pH 7.5 and pH 8.5)
20. 3 M Sodium Acetate (pH 5.2)
21. Dimethyl Sulfoxide
22. 70% ethanol
23. 2 M NaOH
24. Hybridization buffer HT1 (Illumina)

### 2-3. Homemade reagents

Water used should be DNase/RNase-Free Distilled Water

1. 250 mM NaIO_4_: Dissolve 1 mg of Sodium periodate (ACS Reagent Grade) in 18.7 μL of water and keep in the dark. The solution can be aliquoted to 50 μL and stored at −80 °C.
2. 100 mM Biotin (long arm) Hydrazide: Dissolve 50 mg of Biotin (long arm) Hydrazide in 1.345 mL of DMSO. The solution can be aliquoted to 50 μL and stored at −80 °C.
3. LiCl buffer: mix 3.64 mL of 8M Lithium chloride solution, 0.80 mL of 1 M Tris-HCl (pH 7.5), 0.40 mL of 10w/v% Polyoxyethylene (20) Sorbitan Monolaurate Solution, 0.16 mL of 0.5 M EDTA (pH 8.0) and 3.64 mL of water, and store at room temperature.
4. TE wash buffer: Mix 9.12 mL of water, 400 μL of 1 M Tris-HCl (pH 7.5), 400 μL of 10 w/v% Polyoxyethylene (20) Sorbitan Monolaurate Solution and 80 μL of 0.5 M EDTA (pH 8.0), and store at room temperature.
5. Release buffer: Mix 100 μL of RNase ONE 10x Reaction Buffer, 1.0 μL of 10 w/v% Polyoxyethylene (20) Sorbitan Monolaurate Solution and 899 μL of water, and store at room temperature.
6. 1 × TE buffer: Mix 500 μL of 1 M Tris-HCl (pH 8.0), 100 μL of 0.5 M EDTA (pH 8.0) and 49.4 mL of water, and store at room temperature.
7. 0.1 M NaCl/TE buffer: Mix 500 μL of 1 M NaCl and 4.5 mL of 1 × TE buffer, and store at room temperature.
8. 2.5 μM 5’linker:

a. Mix 4 μL of 1 mM 5’adaptor GN5, 4 μL of 1 mM 5’adaptor down, 4 μL of 1 M NaCl and 28 μL of 1 × TE buffer, and carry out the annealing reaction to generate 100 μM GN5 linker <Annealing reaction> 95 °C, 5 min: − 0.1 °C sec −1 down to 83 °C; 5 min at 83 °C; − 0.1 °C sec−1 down to 71 °C; 5 min at 71 °C; − 0.1 °C sec−1 down to 59 °C; 5 min at 59 °C; −0.1 °C sec−1, to 47 °C; 5 min at 47 °C; −0.1 °C sec−1, to 35 °C; 5 min at 35 °C; −0.1 °C s−1 to 23 °C; 5 min at 23 °C; −0.1 °C sec−1 to 11 °C, and then hold at 11 °C.
b. Mix 1 μL of 1 mM 5’adaptor N6, 1 μL of 1 mM 5’adaptor down, 1 μL of 1M NaCl and 7 μL of 1 × TE buffer and repeat the annealing reaction at step 8.a. to generate 100 μM N6 linker
c. Mix 40 μL of 100 μM GN5 linker from step 8.a. and 10 μL of 100 μM N6 linker from step b.
d. Dilute 100 μM of mixed 5’linkers to 2.5 μM in 0.1 M NaCl/TE buffer. For instance, add 2.5 μL of 100 μM mixed linkers from step c to 97.5 μL of 0.1 M NaCl/TE buffer for 25 samples. 2.5 μM and 100 M mixed 5’linkers can be stored at −20 °C.
9. 2.5 μM 3’ linker:

a. Mix 1 μL of 1 mM 3’adaptor up, 1 μL of 1 mM 3’adaptor down, 1 μL of 1 M NaCl and 7 μL of 1 x TE buffer, and carry out the annealing reaction at the step 8.a. to generate 100 μM of 3’ linker.
b. Dilute 100 μM mixed linker from step 9.a. to 2.5 μM with 0.1 M NaCl/TE buffer (see details at the step 8.d.).

### 2-4. Primers and Linker sequences are listed in Table 1 – 3

**Table 1.**
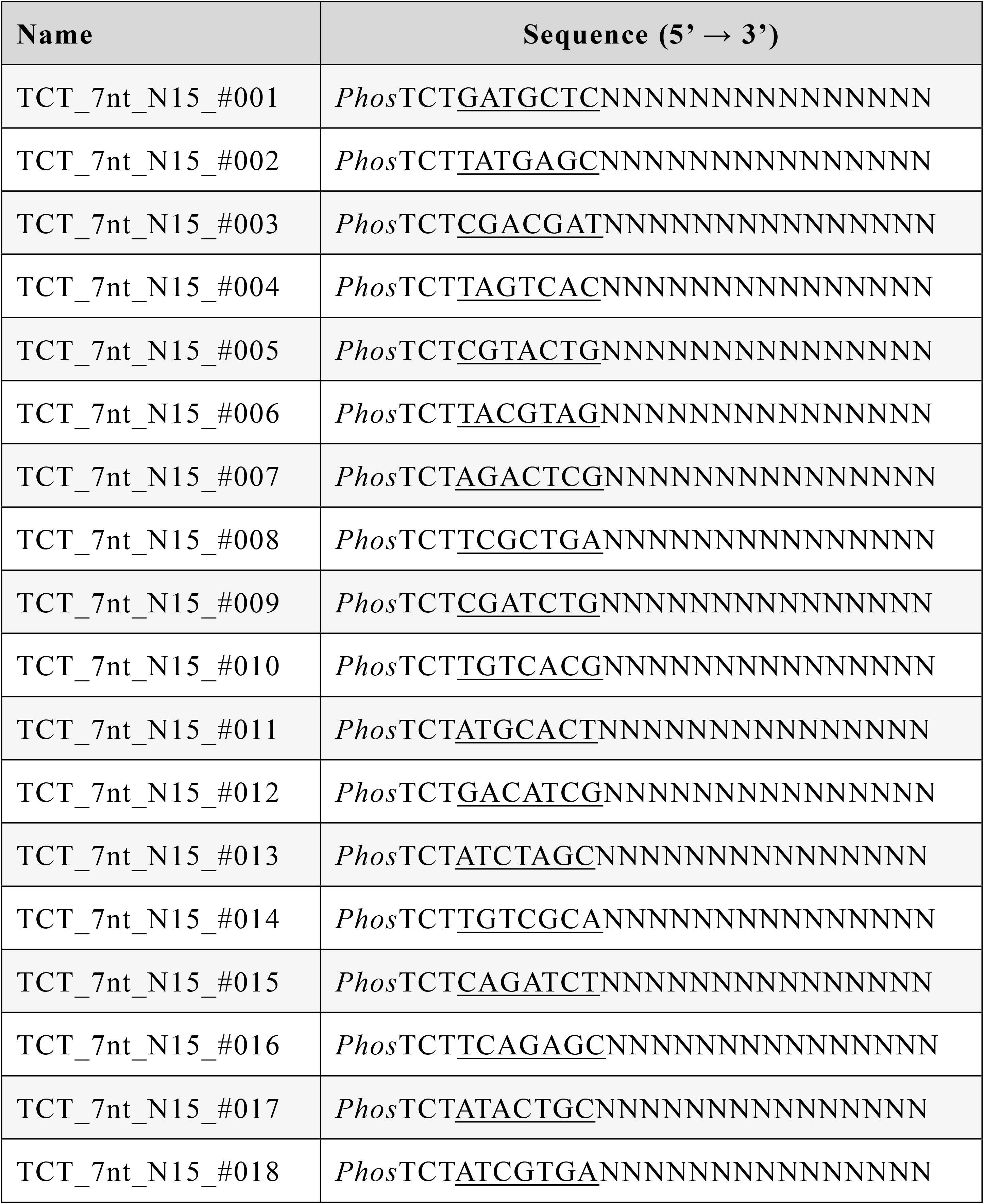
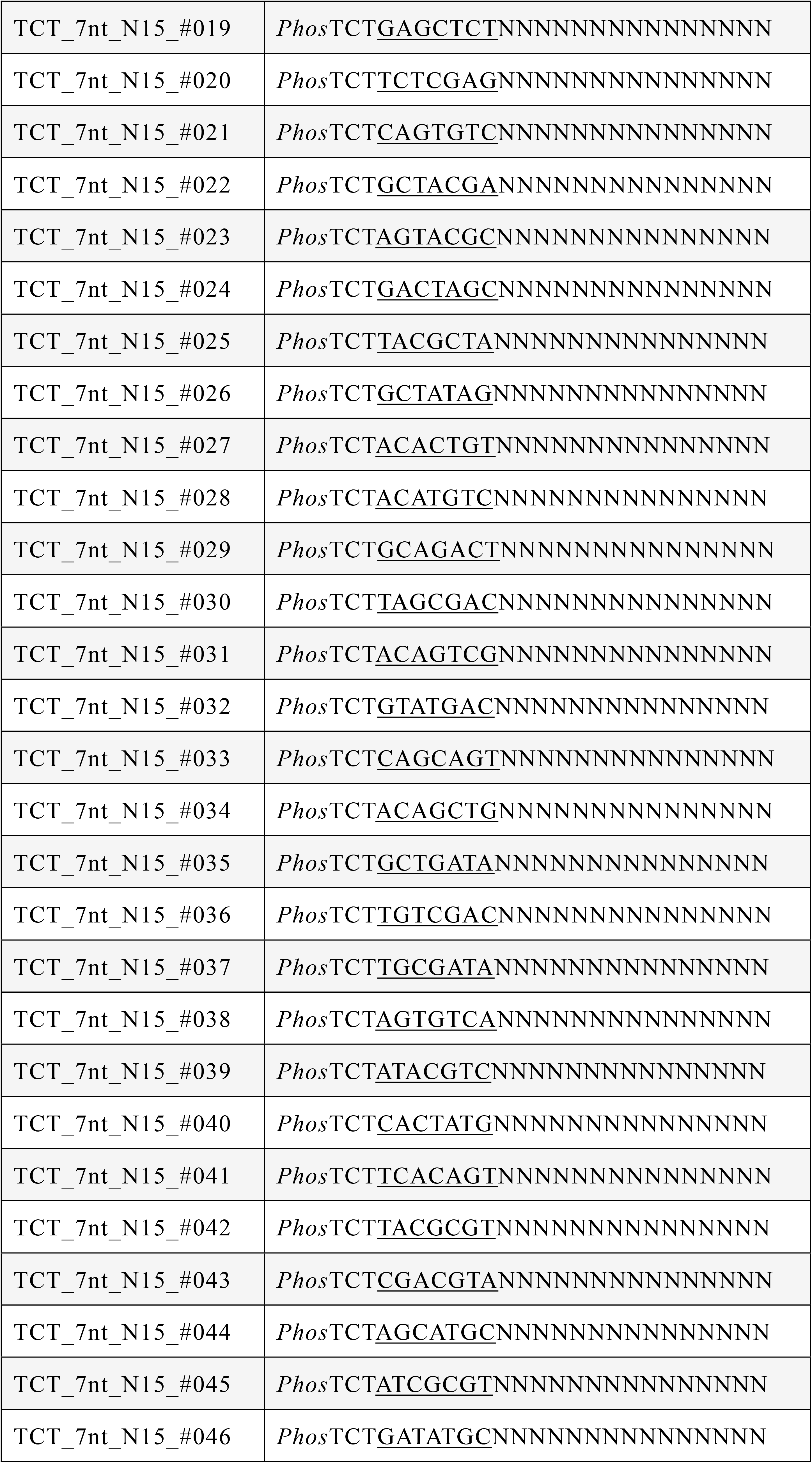
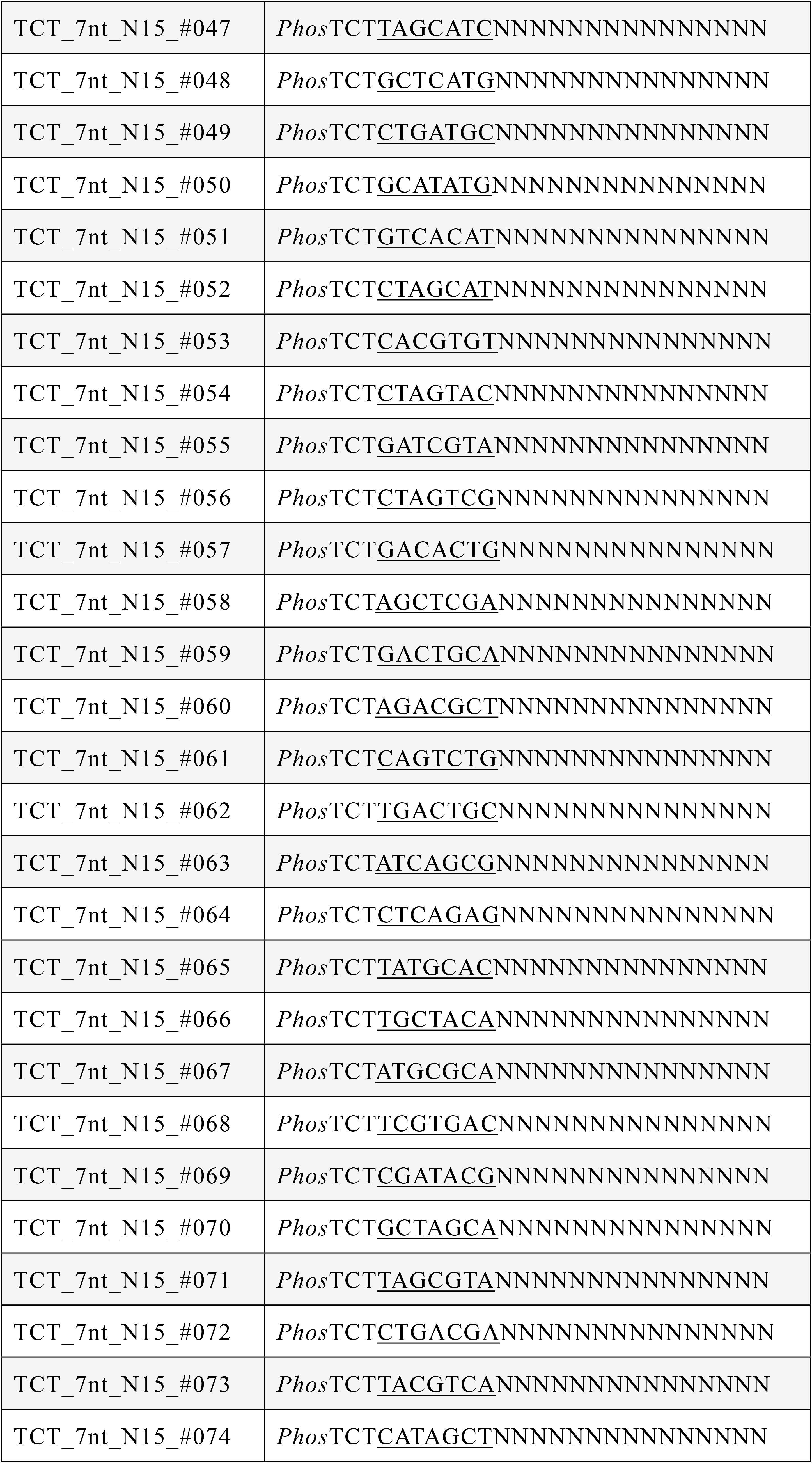
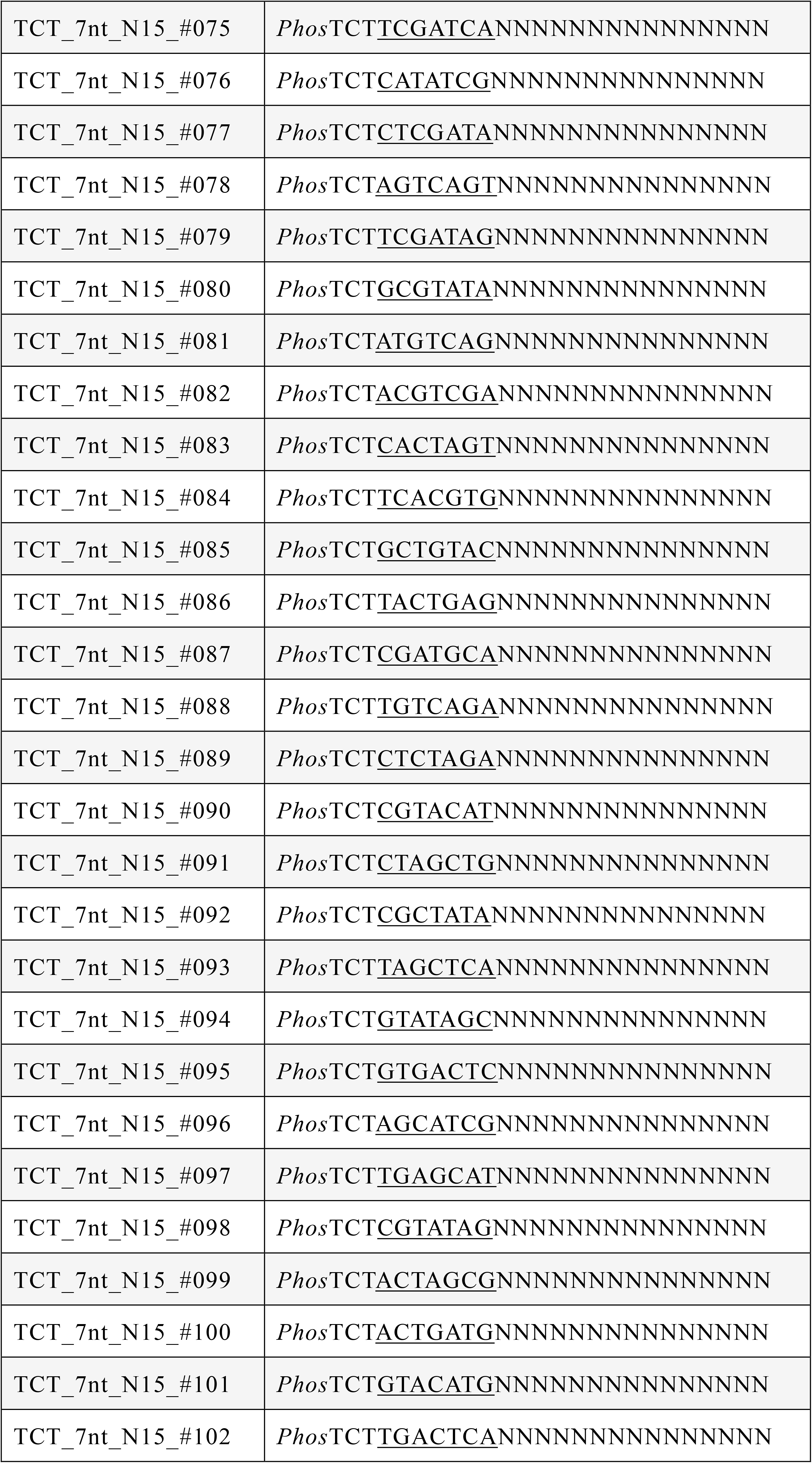
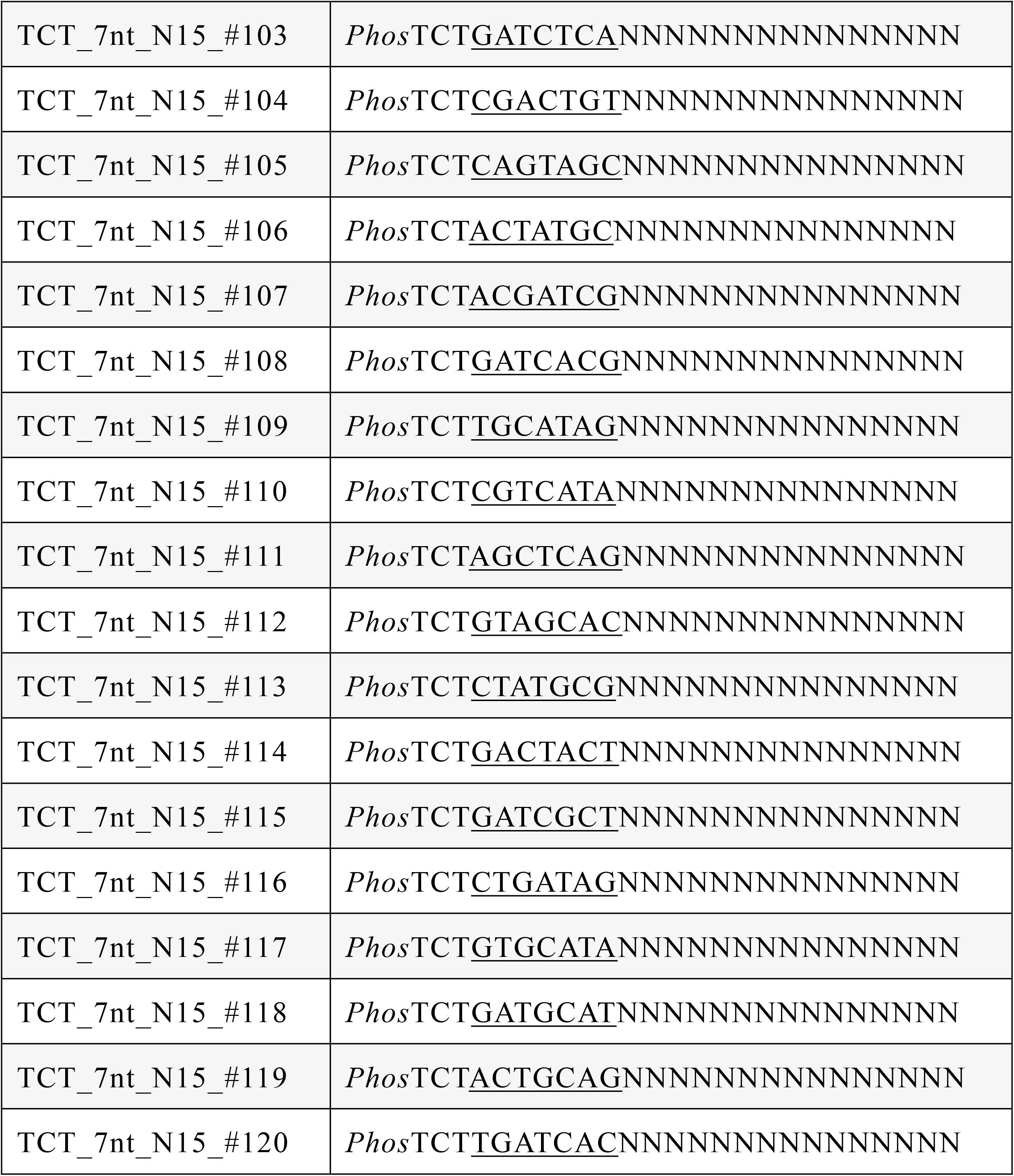
List of RT primers containing barcode

**Table 2.**
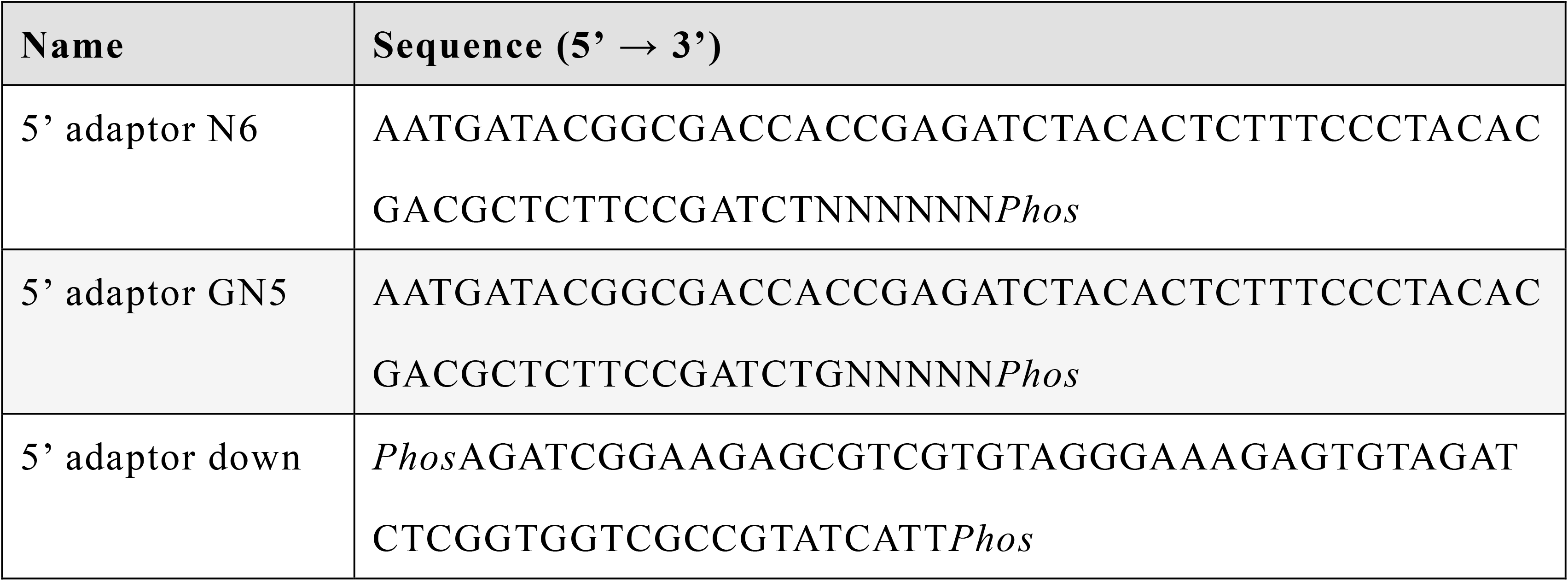
List of 5’ linkers

## 3. Methods (protocol)

Water used should be DNase/RNase-Free Distilled Water

### 3.1 Reverse Transcription (Timing: 5.5 h)

1. Mix 4 μL of 50 ng RNAs (12.5 ng/μL) and 1 μL of 1.25 mM RT primers in a 96 wells plate by pipetting to generate RNA and primer mix on ice (*see* **Note 1 and 2**).
2. Incubate the RNA-primer mix form step 3.1.1 at 65 °C for 5 min and immediately put on ice for 2 min.
3. Mix the following components (enzyme mix) (*see* **Note 3**)

4. Add 5 μL of enzyme mix from step 3 to RNA-primer mix solution from step 3.1.2 and carefully mix 10 times by pipetting on ice.
5. Incubate at 25 °C for 30 sec, 50 °C for 30 min and keep at 4 °C to generate RNA-cDNA hybrids.
6. Mix samples using following steps (*see* **Note 4**).

| Components of enzyme mix | Volume (μL) | Final concentration |
| --- | --- | --- |
| 5 × First Strand buffer | 2 | 1 × |
| 0.1 M DTT | 0.5 | 0.01 M |
| 10 mM dNTPs | 0.5 | 1 mM |
| RNase free water | 1 | - |
| SuperScript III Reverse Transcriptase | 1 | 200 U |
| Total volume | 5 |  |

①. Transfer each 10 μL of RNA-cDNA hybrids from step 3.1.5 to one of new 1.5 mL tube on ice (total volume is 480 μL).
②. Add 15 μL of water to the first 8 wells at the 96 wells plate from step ①, wash wells by pipetting, and transfer the 15 μL of the solutions to the next 8 wells.
③. Wash the 8 wells by pipetting and transfer 15 μL of the solutions to the next 8 wells.
④. Repeat step ③ three times until the end of 8 wells and transfer all solutions (total volume is 120 μL (15 μL × 8 wells) to the 1.5 mL tube from step ① (final volume is 600 μL).
⑤. Mix 600 μL of the solution from step ④ by vortex, spin down and aliquot 200 μL in new two tubes of 1.5 mL tube on ice (total 200 μL in 3 tubes of 1.5 mL tube).
7. Add 300 μL (1.8 -folds) of RNAClean XP beads to the 48 mixed RNA-cDNA hybrids in a 1.5 mL tube from step 3.1.6, mix well by pipetting and then elute the mixed RNA-cDNA hybrids in the following steps at room temperature unless otherwise specified.

①. Incubate for 10 min, spin down and set the 1.5 mL tube on magnetic stand for 5 min.
②. Discard the supernatant by pipette aspiration.
③. Wash the beads with 1.2 mL of 70% ethanol.
④. Set the 1.5 mL tube on magnetic stand for 5 min.
⑤. Discard the supernatant by pipette aspiration.
⑥. Repeat step ③ ~ ⑤ twice.
⑦. Discard the 70% ethanol completely by pipette aspiration.
⑧. Add 100 μL of water and mix by pipetting extensively (more than 60 times) to elute RNA-cDNA hybrids.
⑨. Incubate at room temperature for 5 min.
⑩. Spin down and set the tube on magnetic stand for 5min.
⑪. Transfer 100 μL of the RNA-cDNA hybrids to new 1.5 mL tube.
⑫. Repeat step ⑧ ~ ⑪ twice (final volume is 200 μL).
⑬. Concentrate 200 μL of the RNA-cDNA hybrids solutions from step ⑫ to around 40 μL by SpeedVac vacuum concentrator at 37 °C, and collect the solutions from 3 of 1.5 mL tubes to 1 of 1.5 mL tube (total volume is around 120 μL) and concentrate to 40 μL in the 1.5 mL tube at 37 °C. The timing is around 2 h (*see* **Note 5**).
⑭. Check the sample volume several times during the concentration and adjust the final volume to 40 μL with water when the volume become less than 40 μL.

### 3.2 Oxidation to modify diol group of cap structure (Timing: 10 min)

1. Mix 40 μL of RNA-cDNA hybrid from step 3.1, 2 μL of 1 M NaOAc (pH 4.5) and 2 μL of 250 mM NaIO_4_ by 10 times pipetting on ice.
2. Incubate for 5 min on ice in dark by aluminum foil wrapping.
3. Add 16 μL of 1 M Tris-HCl (pH 8.5) to neutralize the solution and mix well by pipetting on ice.

### 3.3 Purification (Timing: 1 h)

Add 108 μL (1.8 -folds) of RNACleanXP beads to 60 μL of oxidated RNA-cDNA hybrid from step 3.2, mix well by pipetting and then elute the 48 samples mixed RNA-cDNA hybrids in the following steps at room temperature.

1. Incubate for 5min.
2. Spin down and set the tube on magnetic bar for 5 min.
3. Discard the supernatant by pipette aspiration.
4. Wash the beads with 200 μL of 70% ethanol.
5. Discard the 70% ethanol.
6. Repeat step 4 ~ 5 twice and discarded 70% ethanol completely.
7. Add 42 μL of water and mix by pipetting extensively (more than 60 times) to elute supernatant.
8. Incubate for 5min.
9. Spin down and set the tube on magnetic bar for 5 min.
10. Collect 40 μL of the supernatant to new tube.

### 3.4 Biotinylation by the coupling reaction to the oxidized RNA-cDNA hybrids (*see* Note 6) (Timing: 1.5 h)

1. Mix 40 μL of purified oxidized RNA/cDNA hybrid from step 3.3, 4 μL of 1M NaOAc (pH 6.3) and 4 μL of 100 mM Biotin (long arm) hydrazide by 10 times pipetting.
2. Incubate for 30 min at 40 °C.
3. Add 86.4 μL of RNACleanXP (1.8 -folds) to 48 μL of solution from step 3.4.2 and perform purification by following previous step 3.3.

### 3.5 RNaseONE treatment to digest RNA of RNA-cDNA hybrids (*see* Note 7) (Timing: 1.5 h)

1. Add 4.5 μL of 10 × RNaseONE buffer and 0.5 μL of RNaseONE to 40 μL of purified biotinylated RNA-cDNA hybrids from step 3.4 and mix by 10 times pipetting.
2. Incubate for 30 min at 37 °C.
3. Add 81 μL of RNACleanXP (1.8 -folds) to the solution from step 3.5.2 and perform purification by following previous step 3.3.

### 3.6 Dynabeads M-270 Streptavidin beads preparation (Timing: 0.5 h)

1. Add 30 μL of Dynabeads M-270 Streptavidin to new 1.5 mL tube, set on the magnetic stand for 5 min, and discard supernatant.
2. Wash the beads with 30 μL of LiCl buffer, set on the magnetic stand for 5 min, and discard supernatant.
3. Repeat step 3.6.2 twice.
4. Resuspend the beads in 35 μL of LiCl buffer.

### 3.7 CapTrap reaction (Timing: 1 h)

1. Add 35 μL of beads from step 3.6 to 40 μL of RNA-cDNA hybrids from step 3.5 and mix well by pipetting.
2. Incubate for 15 minutes at 37 °C and mix by pipetting at the time of 7 min.
3. Spin down and set the plate on the magnetic bar for 2 min and discard the supernatant by the pipette aspiration.
4. Add 150 μL of TE wash buffer and mix by 60 times pipetting. Spin down and set the plate on the magnetic bar for 2 min and discard the supernatant.
5. Repeat step 4 three times.
6. Add 35 μL of Release buffer to the beads and mix by 60 times pipetting.
7. Incubate for 5 min at 95 °C and on ice for 1 min.
8. Spin down and set the tube on the magnetic bar for 2 min.
9. Transfer the 35 μL of supernatant to new 16 well plate.
10. Add 30 μL of Release buffer to the beads at 3.7.9. and mix well by pipetting, spin down, and set the tube on the magnetic bar for 2 min.
11. Transfer the 30 μL of supernatant to the 16 well plate containing CapTrapped cDNA at step 3.9.9 (total 65 μL).

### 3.8 RNaseONE and RNaseH reaction to remaining RNAs (Timing: 2.5 h)

1. Add 2.9 μL of Release buffer, 2 μL of RNaseONE and 0.1 μL of RNase H to 65 μL of CapTrapped cDNA from step 3.7 and mix by 10 times pipetting.
2. Incubate at 37 °C for 30 min.
3. Add 126 μL of AMPureXP (1.8 -folds) to 33 μL of cDNA from step 3.8.2 and perform purification by following previous step 3.3.
4. Dry up 40 μL of purified CapTrapped cDNA by the SpeedVac concentrator at 37 °C for around 75 min.
5. Add 4 μL of water to the dried pellet.

### 3.9 5’ Single Strand Linker Ligation (Timing: 16.5 h)

1. Incubate 4 μL of CapTrapped cDNA from 3.8 at 95 °C for 5 min and put on ice for 2 min.
2. Incubate 4 μL of 5’ linker at 55 °C for 5 min and put on ice for 2 min.
3. Mix 4 μL of CapTrapped cDNA from step 3.9.1, 4 μL of 5’ linker from step 3.9.2 and 16 μL of Mighty Mix by pipetting (total volume is 24 μL).
4. Incubate at 16 °C for 16 h (overnight).

### 3.10 Purification to remove excess 5’ linkers and linker dimers (Timing: 2 h)

1. Add 46 μL of water and 126 μL of AMPureXP (1.8 -folds) to 24 μL of cDNA from step 3.9 and mix well by pipetting.
2. Perform purification by following previous step 3.3.
3. Incubate at 95 °C for 5 min and immediately put on ice for 2 min.
4. Add 48 μL of AMPureXP beads (1.2 -folds) to 40 μL of cDNA and mix well by pipetting.
5. Perform purification again by following previous step 3.3.
6. Dry up 40 μL of cDNA from step 3.10.5 by SpeedVac concentrator at 80 °C for around 35 min.
7. Add 4 μL of water to the dried pellet.

### 3.11 3’ Single Strand Linker Ligation (*see* Note 8) (Timing: 4.5 h)

1. Incubate 4 μL of cDNA from 3.10 at 95 °C for 5 min and put on ice for 2 min.
2. Incubate 4 μL of 3’ linker at 65 °C for 5 min and put on ice for 2 min.
3. Mix 4 μL of cDNA from step 3.11.1, 4 μL of 3’linker from step 3.11.2 and 16 μL of Mighty Mix by pipetting (total volume is 24 μL).
4. Incubate at 30 °C for 4 h.

### 3.12 Purification to remove excess 3’ linkers and linker dimers (final library) (Timing: 2 h)

1. Purify the solution from step 3.11 by following previous step 3.10 1~6.
2. Add 10 μL of water to the dried pellet.

### 3.13 Library quantification check (Timing: 2.5 h)

Quantify the concentration of library by KAPA Library Quantification Kit in accordance with manufacture’s protocol.

### 3.14 Sequencing

1. Mix the following components

2. Incubate at room temperature for 5 min.
3. Put the tube on ice and add 20 μL of 1 M Tris-HCl (pH 7.0, pre-chilled) to neutralize.
4. Add 110 μL of HT1 buffer.
5. Transfer 120 μL of the denatured and diluted library to HiSeq2500 with 50 bp Paired-End sequencing (see Fig. 1 for the library structure). Read 1 is to read sequence information of cDNA, read 2 is to read sequence information of barcode in RT primer and index read 1 is to read sequence information of index at the 3’ linker. Read 1 contains cDNA sequence, read 2 contains barcode sequence of the RT primer and index of read 1 contains sequence of the index at the 3’ linker.

| Components | Volume (μL) |
| --- | --- |
| Library (2250 amol) | X |
| water | 19-X |
| 2N NaOH | 1 |
| Total volume | 20 |

### 3.15 CAGE tags mapping and gene expression quantification

CAGE tags were mapped to the human genome assembly hg38 using STAR (version 2.5.3a). The average mapping rate was 69.5%, with ~500,000 mapped counts obtained on average across all 240 samples. First, expression for CAGE promoters was estimated by counting the numbers of mapped CAGE tags falling under the 379,952 promoter regions of FANTOM 6 CAT gene models (described in [19]). Next, the expression of the corresponding 124,047 genes was estimated by summing up the expression values of all promoters assigned to a given gene. We found 54,100 genes to have at least one CAGE tag across all 240 libraries. CAGE sequencing summary, raw and summarized gene expression tables are available at https://fantom.gsc.riken.jp/6/suppl/Takahashi_et_al_2020/.

### 3.16 CAGE library correlations

Expression values for the 54,100 genes with at least one CAGE tag across all 240 libraries were correlated for all pairs of CAGE libraries using the Pearson correlation from the ‘cor’ function (‘*stats*’ R package 4.0.0). Correlation for four libraries chosen from two different sequencing runs and with two different barcodes were plotted using ‘plotCorrelation2’ function [20] with ‘tagCountThreshold = 1’ and ‘applyThresholdBoth = FALSE’ parameters.

### 3.17 Genomic and epigenomic gene classes

DHS_type, genomic and epigenomic classifications of genes were inherited from FANTOM CAT annotations [13] and the numbers were plotted for all genes with at least one CAGE tag across all 240 libraries and with available annotations. Genomic classes: “short_ncRNA”, “uncertain_coding”, “small_RNA” and “structural_RNA” were broadly classified as ‘other’. Detailed annotations of all genes are available at https://fantom.gsc.riken.jp/6/suppl/Takahashi_et_al_2020/.

## 4. Results

### Sequencing coverage and reproducibility

We made 240 samples from 50 ng of RNA extracted from THP-1 cells and generated a total of ~134.44 million reads (median of 575,008 reads) using paired end 50 cycles kit on Illumina HiSeq-2000 sequencers in two independents sequencing runs. Reproducibility across all biological replicates was generally quite high (Pearson’s correlation coefficients 0.93-0.98). Examples of correlations for four LQ-ssCAGE libraries with different barcodes selected from two different sequencing runs are shown in Fig. 2.

**Fig. 2.**
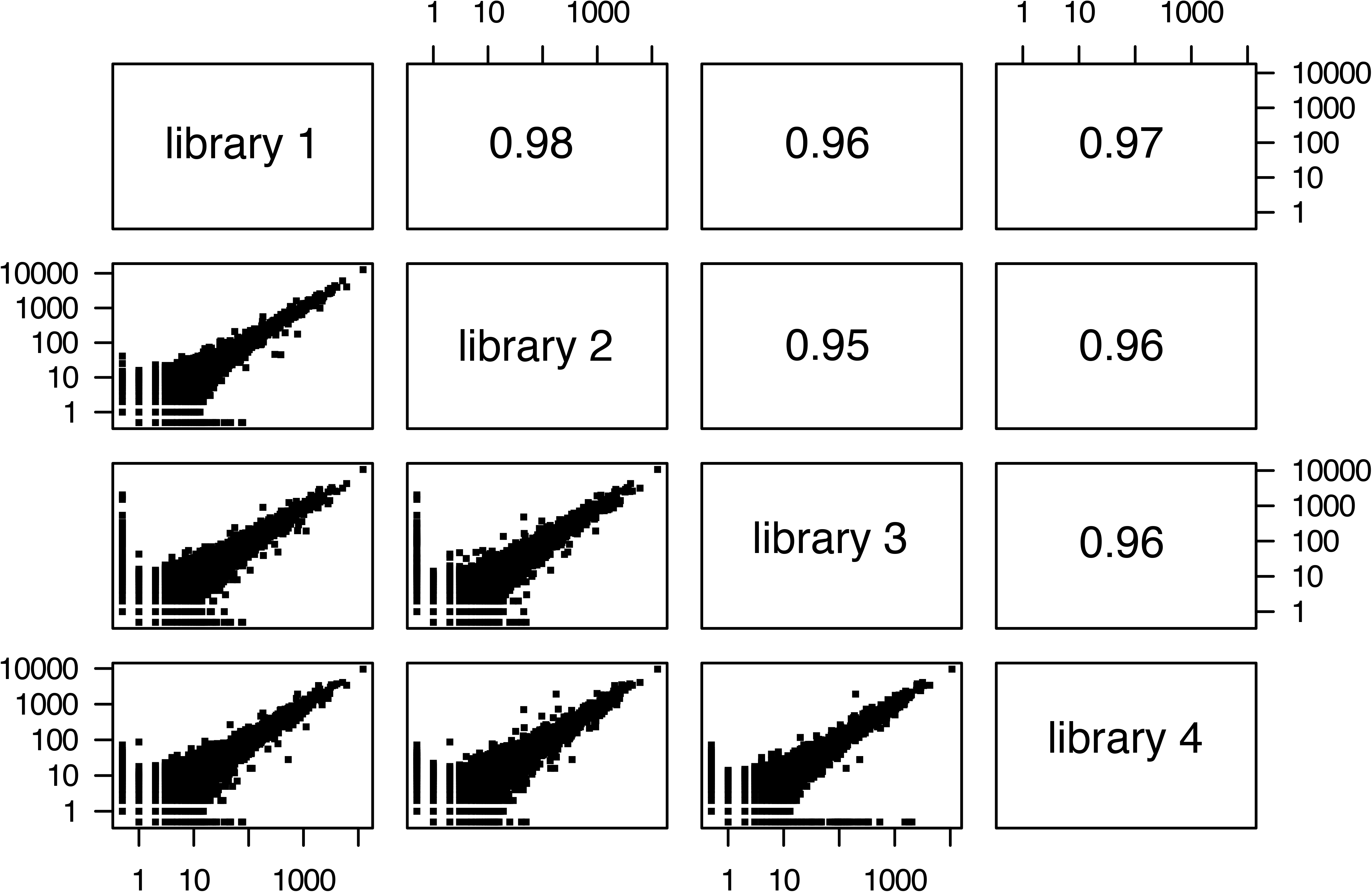
Person correlations between four selected biological replicates in THP-1 whole cell showing technical reproducibility of the method.

### Annotated genes coverage

Using the FANTOM CAT gene annotations [13] we cover (defined as identification of at least one CAGE tag in at least one library) for 54,100 out of 124,047 genes/transcripts, ranging from 4,403 to 18,616 for sense-overlap-RNAs and protein-coding genes, respectively (Fig. 3a). From these, 6,920 were annotated as enhancer (e) lncRNA, 5,817 as promoter-derived intergenic lncRNA and 1,331 as promoter-derived divergent lncRNA (Fig. 3b), when considering a subset of 44,069 genes with their DHSs classified as either enhancer or promoter [18]. Specifically, LQ-ssCAGE method has an advantage of capturing many lowly expressed e-lncRNAs (Fig. 3c): we found 6,920 of all e-lncRNAs in one cellular state, which is definitely a large number considering the high cell-type specificity of e-lncRNAs expression. Selected examples, a novel e-lncRNA (*CATG00000112934.1*) and a p-lncRNA (*GAS5*) with the average expression of ~22 transcripts per million (TPM) and ~226 TPM, respectively, are shown in Fig. 4.

**Fig. 3.**
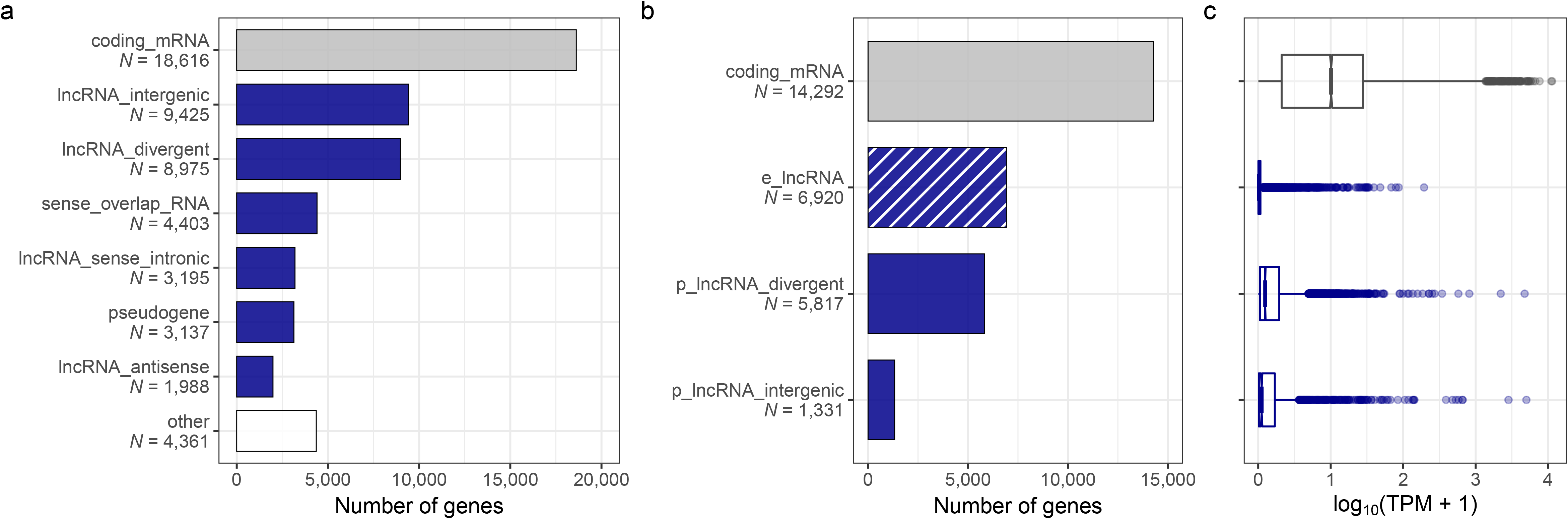
Coverage and expression levels of annotated genes in LQ-ssCAGE: a) number of genes in a given gene classes with at least one CAGE tag in one library as defined in [13], b) number of genes with DNase I hypersensitive sites (DHSs) classified as either enhancer or promoter [18], c) mean expression levels of each epigenetic gene class as on b.

**Fig. 4.**
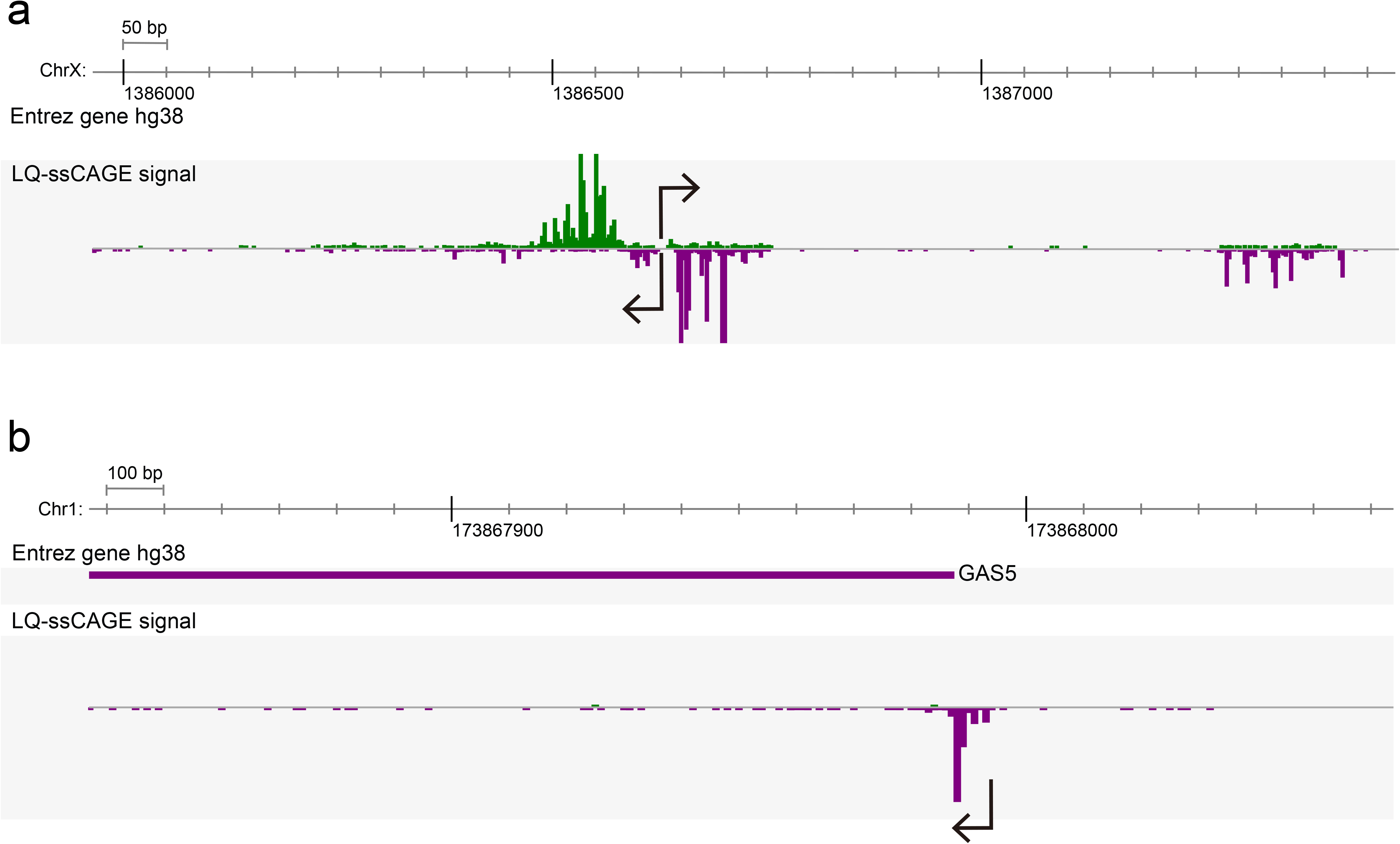
Genome browser views of selected examples of a) e-lncRNA (novel gene) and b) p-intergenic-lncRNA (GAS5). CAGE tags in green are corresponded to the sense strand and in violet are corresponded to the antisense strand.

## 5. Discussion

Since its first publication in 2003 using sanger sequencing, the family of CAGE protocols has evolved to accommodate higher throughput, better cost performance and deeper sequencing, thanks also to the appearance of next generation sequencing [1–6]. Here, we report and protocol that enable expression profiling of large, scalable number of samples from small number of cells in a 96 well plate format. The method enables profiling gene expression with simultaneous promoter and enhancer maps, as well as broad measurement of expression of non-coding, capped RNAs. The protocol is scalable to a large number of plates to serve high throughput screening of parallel assays, including perturbation on large scale on small amounts of cells. Because LQ-ssCAGE does not use PCR, the presented protocol is ideal to measure fine differences of gene expression among different conditions, providing a solid solution to expression profiling for assays to test function of genes, promoters, lncRNAs and screening of molecules that alter cell behavior.

## Notes

1. The amount of RNAs in each tube should be 25 ng ~ 100 ng. After the reverse transcription, samples will be barcodes and can be mixed, using up to 5 μg/tube. For instance, if the starting RNA amount is 50 ng, 100 samples can be mixed in one tube for subsequence analysis. If starting from 100ng of RNAs, process the pooled library in two tubes. The number of mixed samples is dependent on how much samples an operator needs to perform. We describe here, for example, a mix of 48 samples from 50 ng starting RNA. Users can modify these numbers with taking care of key points (see below).
2. The mixed solution should contain no more than 5 μg of starting sample, otherwise the following linkers at step 3.9 and 3.11 may become insufficient for the linker ligation.
3. We advise to prepare premix solution for the sample numbers times 1.1 the number of samples.
4. Step 3.1.6 is for the collection of remaining molecules to avoid losing any RNA-cDNA hybrids in the wells. We recommend to aliquot less than 200 μL per each 1.5 mL tube at step 3.1.6. ⑤ due to the volume limitations of next RNAClean XP purification step.
5. **DO NOT DRY UP** the RNA-cDNA hybrids solutions by the concentrating step because RNAs stick very easily to the surface of tubes when dried.
6. Biotinylation occurs at both 5’end of capped RNAs and 3’end of RNA (described in Fig. 2 in the published protocol [4]), but RNAse treatment removes the 3’ end biotin.
7. RNase treatment is critical to digest uncompleted cDNA synthesis to the 5’ end of capped RNAs (described in Fig. 2 at published protocol [4]).
8. When generating more than 2 libraries with same barcode in the RT primers, use different Index in the 3’linker to mix (*see* Table 3).

**Table 3.**
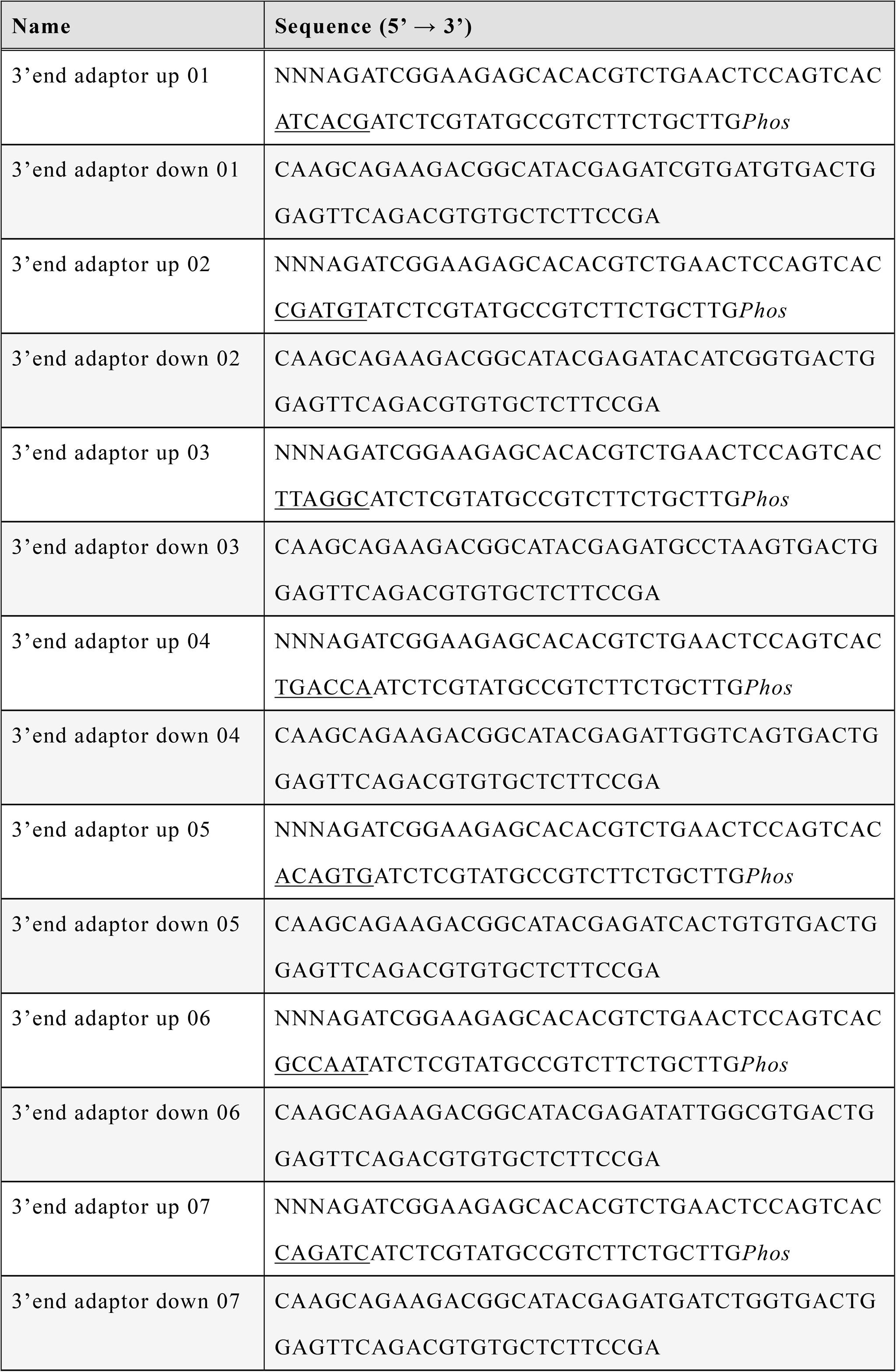
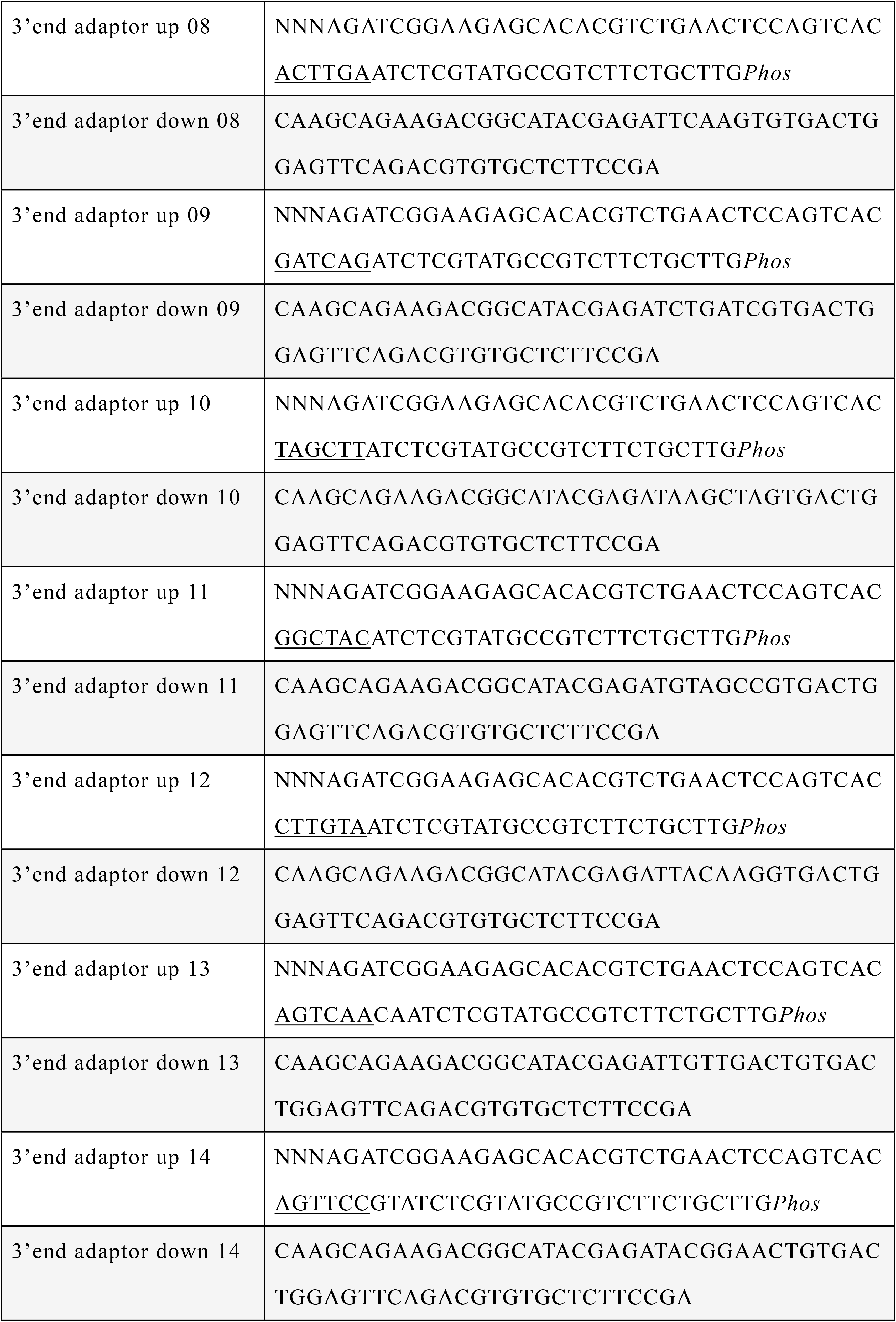
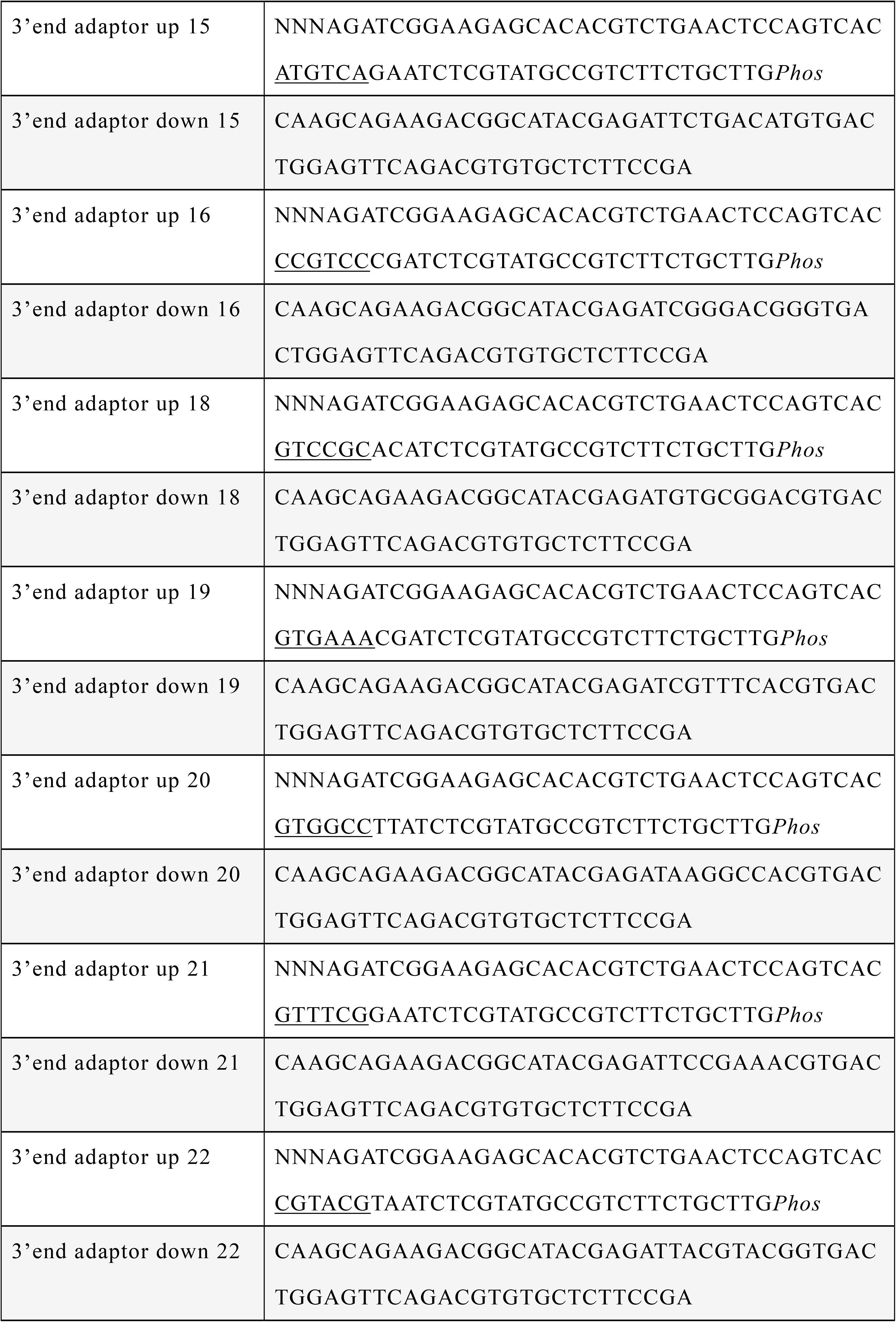
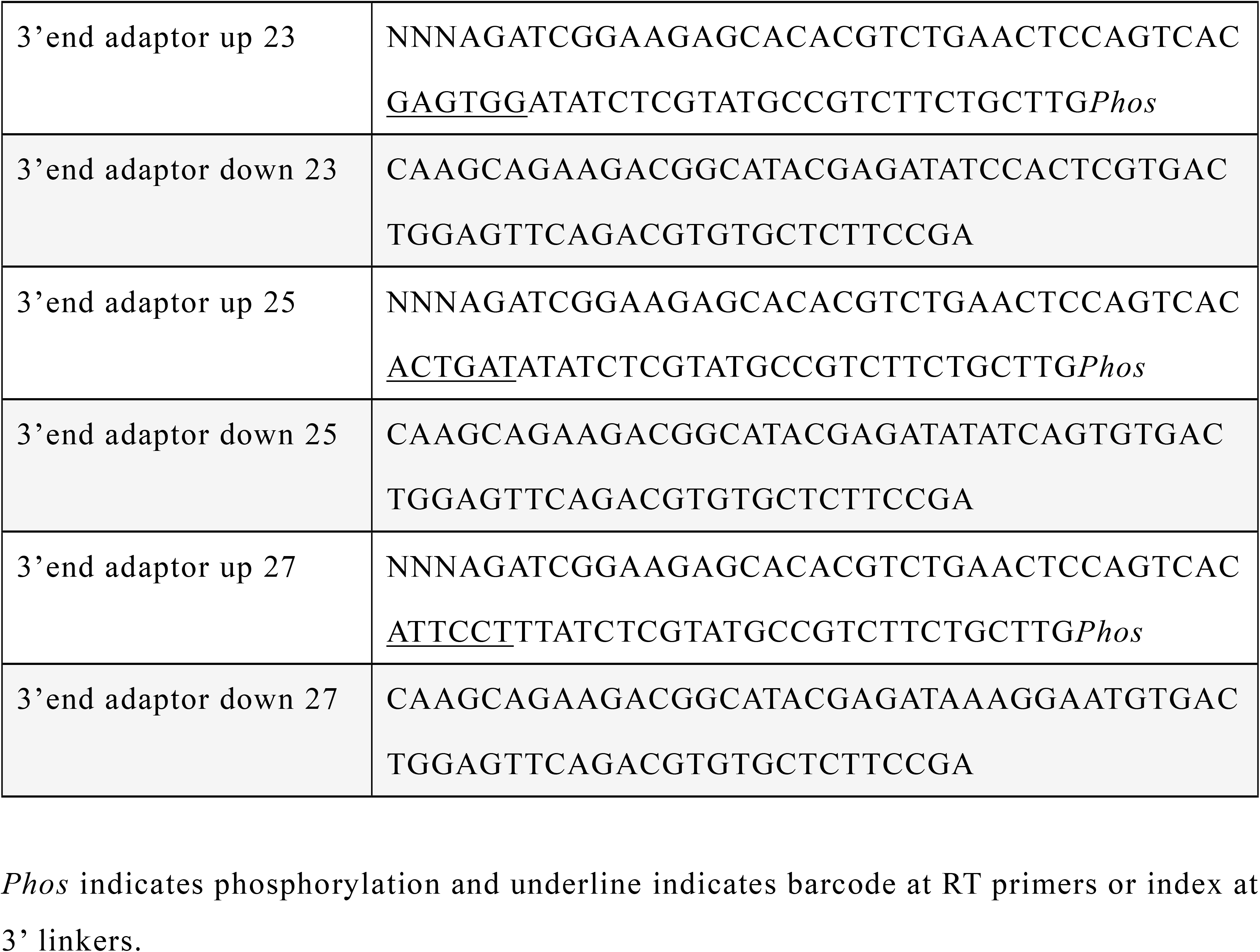
List of 3’ linkers

## Acknowledgement

We special thank our colleagues Mitsuyoshi Murata, Shohei Noma, Miki Kojima and Michihira Tagami for the technical development and Laboratory for Comprehensive Genomic Analysis IMS RIKEN for the sequencing service. This work was supported by a Research Grant from the Japanese Ministry of Education Culture, Sports, Science and Technology (MEXT) to the RIKEN Center for Integrative Medical Sciences.

## References

1. Shiraki T, Kondo S, Katayama S, Waki K, Kasukawa T, Kawaji H, et al. Cap analysis gene expression for high-throughput analysis of transcriptional starting point and identification of promoter usage. Proc Natl Acad Sci U S A. 2003;100(26):15776–81. doi: 10.1073/pnas.2136655100.

2. Kodzius R, Kojima M, Nishiyori H, Nakamura M, Fukuda S, Tagami M, et al. CAGE: cap analysis of gene expression. Nat Methods. 2006;3(3):211–22. doi: 10.1038/nmeth0306-211.

3. Plessy C, Bertin N, Takahashi H, Simone R, Salimullah M, Lassmann T, et al. Linking promoters to functional transcripts in small samples with nanoCAGE and CAGEscan. Nat Methods. 2010;7(7):528–34. doi: 10.1038/nmeth.1470.

4. Takahashi H, Lassmann T, Murata M, Carninci P. 5 ’ end-centered expression profiling using cap-analysis gene expression and next-generation sequencing. Nature Protocols. 2012;7(3):542–61. doi: 10.1038/nprot.2012.005.

5. Murata M, Nishiyori-Sueki H, Kojima-Ishiyama M, Carninci P, Hayashizaki Y, Itoh M. Detecting expressed genes using CAGE. Methods Mol Biol. 2014;1164:67–85. doi: 10.1007/978-1-4939-0805-9_7.

6. Cvetesic N, Leitch HG, Borkowska M, Müller F, Carninci P, Hajkova P, et al. SLIC-CAGE: high-resolution transcription start site mapping using nanogram-levels of total RNA. Genome Res. 2018;28(12):1943–56. doi: 10.1101/gr.235937.118.

7. Andersson R, Sandelin A. Determinants of enhancer and promoter activities of regulatory elements. Nat Rev Genet. 2020;21(2):71–87. doi: 10.1038/s41576-019-0173-8.

8. Consortium EP. An integrated encyclopedia of DNA elements in the human genome. Nature. 2012;489(7414):57–74. doi: 10.1038/nature11247.

9. Forrest AR, Kawaji H, Rehli M, Baillie JK, de Hoon MJ, Haberle V, et al. A promoter-level mammalian expression atlas. Nature. 2014;507(7493):462–70. doi: 10.1038/nature13182.

10. Andersson R, Gebhard C, Miguel-Escalada I, Hoof I, Bornholdt J, Boyd M, et al. An atlas of active enhancers across human cell types and tissues. Nature. 2014;507(7493):455–61. doi: 10.1038/nature12787.

11. Sanyal A, Lajoie BR, Jain G, Dekker J. The long-range interaction landscape of gene promoters. Nature. 2012;489(7414):109–13. doi: 10.1038/nature11279.

12. Rennie S, Dalby M, van Duin L, Andersson R. Transcriptional decomposition reveals active chromatin architectures and cell specific regulatory interactions. Nat Commun. 2018;9(1):487. doi: 10.1038/s41467-017-02798-1.

13. Hon CC, Ramilowski JA, Harshbarger J, Bertin N, Rackham OJ, Gough J, et al. An atlas of human long non-coding RNAs with accurate 5’ ends. Nature. 2017;543(7644):199–204. doi: 10.1038/nature21374.

14. Haberle V, Li N, Hadzhiev Y, Plessy C, Previti C, Nepal C, et al. Two independent transcription initiation codes overlap on vertebrate core promoters. Nature. 2014;507(7492):381–5. doi: 10.1038/nature12974.

15. Carninci P, Kvam C, Kitamura A, Ohsumi T, Okazaki Y, Itoh M, et al. High-efficiency full-length cDNA cloning by biotinylated CAP trapper. Genomics. 1996;37(3):327–36. doi: 10.1006/geno.1996.0567.

16. Pillay S, Takahashi H, Carninci P, Kenhere A. Antisense ncRNAs during early vertebrate development are divided in groups with distinct features. bioRxiv; 2020.

17. Chen L, Yang W, Guo Y, Chen W, Zheng P, Zeng J, et al. Exosomal lncRNA GAS5 regulates the apoptosis of macrophages and vascular endothelial cells in atherosclerosis. PLoS One. 2017;12(9):e0185406. doi: 10.1371/journal.pone.0185406.

18. Kundaje A, Meuleman W, Ernst J, Bilenky M, Yen A, Heravi-Moussavi A, et al. Integrative analysis of 111 reference human epigenomes. Nature. 2015;518(7539):317–30. doi: 10.1038/nature14248.

19. Ramilowski JA, Yip CW, Agrawal S, Chang JC, Ciani Y, Kulakovskiy IV, et al. Functional annotation of human long noncoding RNAs via molecular phenotyping. Genome Res. 2020. doi: 10.1101/gr.254219.119.

20. Haberle V, Forrest AR, Hayashizaki Y, Carninci P, Lenhard B. CAGEr: precise TSS data retrieval and high-resolution promoterome mining for integrative analyses. Nucleic Acids Res. 2015;43(8):e51. doi: 10.1093/nar/gkv054.

